# Population genetic structure of the bank vole *Myodes glareolus* within its glacial refugium in peninsular Italy

**DOI:** 10.1101/467753

**Authors:** Andrea Chiocchio, Paolo Colangelo, Gaetano Aloise, Gianni Amori, Sandro Bertolino, Roberta Bisconti, Riccardo Castiglia, Daniele Canestrelli

**Affiliations:** Department of Ecological and Biological Science, Università degli Studi della Tuscia, I-01100 Viterbo, Italy; National Research Council, Institute of Research on Terrestrial Ecosystems, Via Salaria km 29.300, I-00015 Monterotondo (Rome), Italy; Museo di Storia Naturale ed Orto Botanico, Università della Calabria. Via Savinio s.n.c., Edificio Polifunzionale, I - 87036 Rende (CS), Italy; Department of Biology and Biotechnology “Charles Darwin”, University of Rome La Sapienza, Italy, I-00185 Roma, Italy; Department of Life Sciences and Systems Biology, Università degli Studi di Torino, I-10123 Torino, Italy

**Keywords:** Keywords:, Italian peninsula, glacial refugia, *Myodes glareolus*, phylogeography, genetic structure

## Abstract

It is now well established that Southern European peninsulas have been major glacial refugia for temperate species during Pleistocene climatic oscillations. However, substantial environmental changes occurred also within these peninsulas throughout the Pleistocene, rising questions about the role and interplay of various of micro-evolutionary processes in shaping patterns of intraspecific diversity within these areas. Here, we investigate the patterns of genetic variation in the bank vole *Myodes glareolus* within the Italian peninsula. By using a panel of 13 microsatellite loci, we found more intra-specific variation than expected based on previous assessments. Indeed, both Bayesian and ordination-based clustering analyses of variation recovered five main geographic/genetic clusters along the peninsula, with three clusters geographically restricted to the southern portion of the study area. This pattern supports the occurrence of multiple sub-refugia for the bank vole in peninsular Italy, likely promoted by the major paleo-environmental changes which affected forested habitats within this area during the Pleistocene. Thus, our results support a scenario whereby the high levels of intraspecific diversity observed within major glacial refugia are better explained by dynamic micro-evolutionary processes occurred within these areas, rather than by long-term demographic stability of refugial population. Finally, the narrow and isolated distribution of some of the identified lineages, suggest the need for future assessments of their conservation and taxonomic status.

## Introduction

Southern European peninsulas have provided an excellent research ground to investigate how past climate changes and topographic features influenced species’ evolutionary histories (Hewitt, 2011). Plenty of studies in the last thirty years highlighted the role of these peninsulas as climatic refugia for temperate species during Pleistocene glacials (Bennet & Provan, 2008; Comes & Kadereit, 1998; Feliner, 2011; Hewitt, 1996; 2004; Schmitt, 2007; Weiss & Ferrand, 2007; Stewart, Lister, Barnes & Dalén, 2010). Due to the strong topographic complexity of these peninsulas, species underwent extreme population fragmentation within refugia (Gomez & Lunt, 2007; Hofreiter & Stewart, 2009). As a consequences, these areas have been found to be particularly rich of intraspecific genetic lineages – sometimes highly divergent from the closest relatives (e.g. Canestrelli, Cimmaruta, Costantini & Nascetti, 2006), often with narrow distribution (e.g. Bisconti et al., 2018) – detecting which is crucial to understanding species genetic structure and to defining evolutionary and management units for conservation planning (Avise, 2008; Frankham, 2010; Groves et al., 2017; Palsbøll, Berube & Allendorf, 2007). However, despite their disproportionate importance for the historical biogeography of Western Palearctic biota, current knowledge of their biodiversity patterns is still far from satisfactory. For many taxa, including charismatic taxa like mammals, the genetic structure is still little known, and the available knowledge if often flawed by limited sampling or limited number of genetic markers.

The bank vole *Myodes glareolus* (Schreber, 1780) is a small woodland-dwelling rodent, widespread throughout temperate and boreal forests of most of Europe (Amori, Contoli & Nappi, 2008a), which has been a key species in the study of the European fauna response to the Pleistocene climate changes. It has been one of the most convincing examples of a woodland species surviving glaciations within a cryptic northern refugium in Europe, i.e. a refugium located further north of the traditionally recognized refugia in the Southern European peninsulas (Bhagwat & Willis, 2008; Bilton, Mirol, Mascheretti, Fredga, Zima & Searle, 1998; Deffontaine et al., 2005; Filipi, Marková, Searle & Kotlík, 2015; Kotlík, Deffontaine, Mascheretti, Zima, Michaux, & Searle, 2006). However, bank vole populations survived Pleistocene glaciations also in southern refugia. There were identified distinct evolutionary lineages in either Balkan, Iberian and Italian peninsulas (Colangelo, Aloise, Franchini, Annesi & Amori, 2012; Deffontaine et al., 2005; Filipi et al., 2015). Within the Italian peninsula, four distinct evolutionary lineages have been characterized by mean of mitochondrial DNA variation: one widespread across Alps and northern Italy, one distributed mainly throughout northern and central Apennines, one restricted to the Gargano promontory (Apulia), and one found only in Calabria (Colangelo et al., 2012). This differentiation is supported also by some slight morphological distinctiveness (Amori et al., 2008a). Interestingly, the Calabrian clade showed strong and ancient (Early Pleistocene) genetic divergence from all other *M. glareolus* lineages, resulting as the basal clade of the entire bank vole phylogeny, whereas the Gargano clade does not cluster with any of the other lineages (Colangelo et al., 2012; Filipi et al., 2015). Nevertheless, all phylogenetic and phylogeographic inferences were based only on mitochondrial data, which have several limitations in inferring population genetic structure and patterns of gene flow among populations (Ballard & Whitlock, 2004). Indeed, despite Calabrian and Apulian populations resulted genetically differentiated from the other peninsular populations, and virtually geographically isolated by the rarefaction/fragmentation of species habitat, the lack of a multi-marker analysis of species genetic structure does not allow inferences on genetic isolation.

In this study, we further investigate the genetic structure of the bank vole in the Italian peninsula. We employ a set of thirteen microsatellite loci in order to complement previously published mitochondrial data (Colangelo et al., 2012), with the aim of better understanding the geographic structure of genetic variation and to shed more light on the bank vole evolutionary history. Moreover, considering the narrow ranges of the Calabrian and Apulian lineages and the ongoing reduction of the forest habitat (Scarascia-Mugnozza, Oswald, Piussi & Radoglou, 2000), a better understanding of the pattern of genetic isolation is mandatory to evaluate the need for conservation actions concerning southern bank vole populations.

## Materials and methods

We collected 76 *Myodes glareolous* individuals from 15 localities spanning the Italian peninsula; collecting sites and sample sizes are given in Table 1 and Figure 1. Tissue samples were obtained from an auricle biopsy on live-trapped animals during field sessions, or from museum specimens (Museum of Comparative Anatomy G.B. Grassi of the University of Rome ‘La Sapienza’).

**Figure 1.**
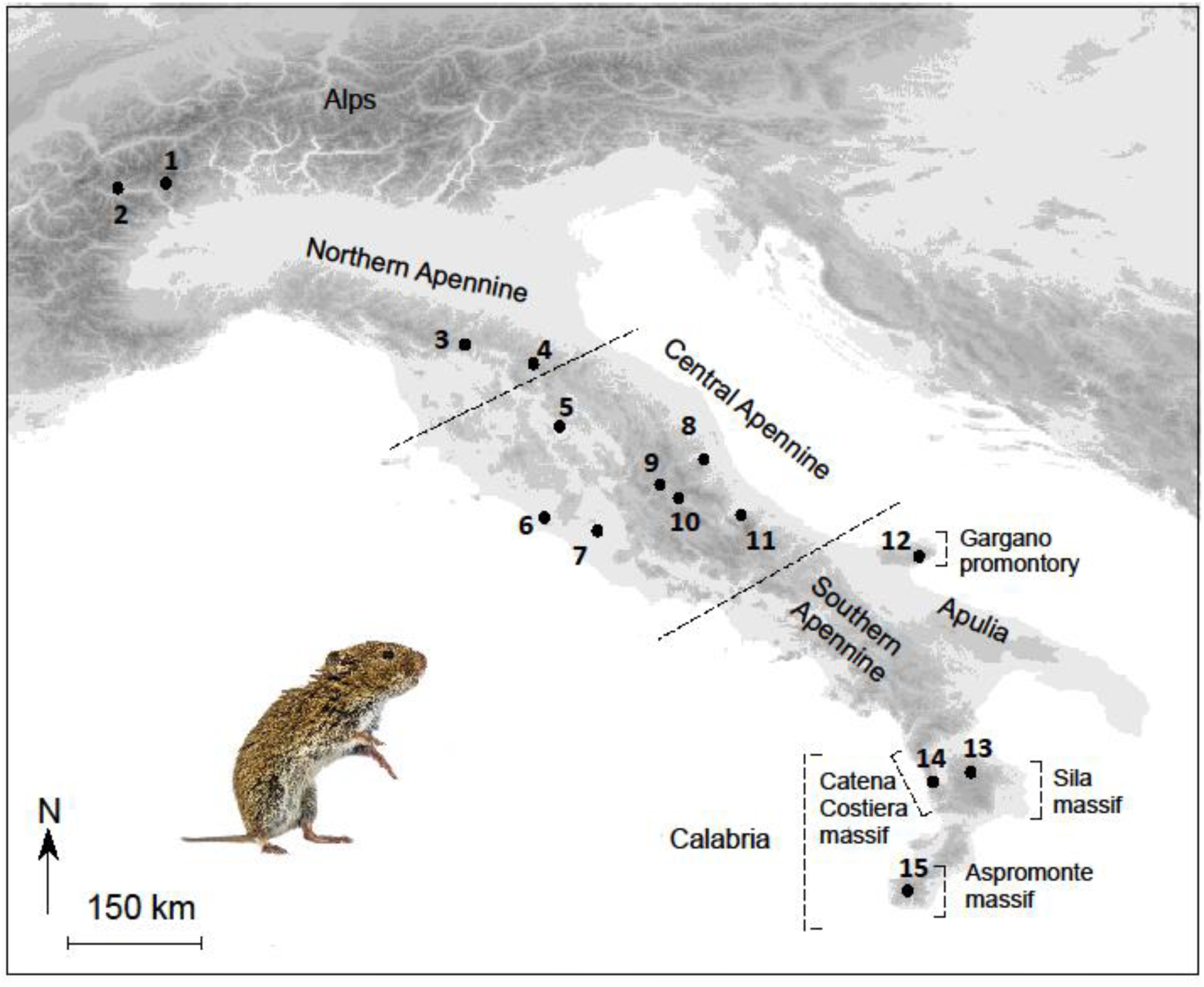
Geographical localization of the 15 populations of *Myodes glareolus* sampled and analysed. Localities are numbered as in Table 1; dashed lines delimits the main geographic regions named in the text. The map was drawn using the software Canvas 11 (ACD Systems of America, Inc.). Photo: *Myodes glareolus* (from Rudmer Zwerver via Photodune.net).

**Table 1.** Sample number, collecting locality, sample size (analyzed specimens), geographic coordinates, allelic richness (Ar), and expected heterozigosity (He).

| Sample | Locality | N | Latitude | Longitude | Ar | He |
| --- | --- | --- | --- | --- | --- | --- |
| 1 | Gressoney | 1 | 45.76902 | 7.82552 | - | - |
| 2 | Aosta | 11 | 45.73644 | 7.31298 | 2.302 | 0.594 |
| 3 | Cantagallo | 1 | 44.03176 | 11.05226 |  | - |
| 4 | Foresta della Lama | 10 | 43.81710 | 11.81270 | 2.869 | 0.718 |
| 5 | Passignano sul Trasimeno | 1 | 43.21675 | 12.13462 | - | - |
| 6 | Tolfa | 1 | 42.14959 | 11.93806 | - | - |
| 7 | Settebagni | 1 | 42.13705 | 12.53497 | - | - |
| 8 | Civitella del Tronto | 1 | 42.79336 | 13.67409 | - | - |
| 9 | Montereale | 3 | 42.49670 | 13.19747 | 2.564 | 0.564 |
| 10 | L'Aquila | 4 | 42.35686 | 13.38911 | 2.435 | 0.555 |
| 11 | Majella Massif | 6 | 42.14480 | 14.07530 | 2.584 | 0.605 |
| 12 | Gargano Promontory | 11 | 41.75540 | 16.01180 | 1.757 | 0.417 |
| 13 | Sila Plateau | 14 | 39.37211 | 16.57618 | 2.687 | 0.724 |
| 14 | Catena Costiera | 3 | 39.27578 | 16.09198 | 2.687 | 0.584 |
| 15 | Aspromonte Massif | 8 | 38.15910 | 15.92060 | 2.357 | 0.599 |
|  |  | 76 |  |  |  |  |

DNA extractions were performed by using the standard cetyltrimethylammonium bromide protocol (Doyle and Doyle, 1987). We analysed genetic variation at thirteen microsatellite loci: *Cg14E1*, *Cg15F7*, *Cg17A7*, *Cg3A8*, *Cg3F12*, *Cg4F9*, *Cg12A7*, *Cg13C12*, *Cg17E9*, *Cg10A11*, *Cg10H1*, *Cg13G2*, *Cg6A1* (Guivier et al., 2011; Rikalainen, Grapputo, Knott, Koskela & Mappes, 2008) following protocols published in Guivier et al. (2011). We chose a subset of available loci after excluding those that exhibited reaction inconsistency in over 30% of the samples analysed. The thirteen loci were assembled in three multiplex as described in Table 2. Forward primers were fluorescently labelled, and PCR products were electrophoresed by Macrogen Inc. on an ABI 3730xl genetic analyser (Applied Biosystems) with a 400-HD-size standard.

**Table 2.** Marker name, repeat motif, and colour dye of the thirteen microsatellite loci assembled in three multiplex.

| Multiplex | Marker | Repeat | Dye |
| --- | --- | --- | --- |
| A | Cg4F9 | (CA)<sub>20</sub> | FAM |
|  | Cg14E1 | (CT)<sub>20</sub> | FAM |
|  | Cg17A7 | (ATGT)<sub>9</sub> | HEX |
|  | Cg15F7 | (CT)<sub>20</sub> | HEX |
|  | Cg3F12 | (GT)<sub>16</sub> | TAMRA |
|  | Cg3A8 | (GT)<sub>21</sub> | TAMRA |
| B | Cg12A7 | (GA)<sub>21</sub> | FAM |
|  | Cg17E9 | (GTAT)<sub>9</sub> | FAM |
|  | Cg13C12 | (CT)<sub>21</sub> | HEX |
| C | Cg13G2 | (GT)<sub>14</sub> | FAM |
|  | Cg10H1 | (GACA)<sub>7</sub> | FAM |
|  | Cg10A11 | (GT)<sub>15</sub> | HEX |
|  | Cg6A1 | (CAT)<sub>18</sub> | HEX |

The microsatellite data were analysed using GeneMapper® 4.1. Micro-Checker 2.2.3 (Van Oosterhout, Hutchinson, Wills & Shipley, 2004) was used to test for null alleles and large-allele dropout influences. Allelic frequencies were computed by using GENETIX 4.05 (Belkhir, Borsa, Chikhi, Raufaste & Bonhomme, 1996), while FSTAT (Goudet, 1995) was used to test for deviations from the expected Hardy-Weinberg and linkage equilibria. Estimates of genetic diversity, based on the mean allelic richness and the mean observed and expected heterozygosity were computed by the DivRsity R package (Keenan, McGinnity, Cross, Crozier & Prodöhl, 2013), after excluding populations with n < 4; allelic richness was computed using the rarefaction method (Petit, Mousadik & Pons, 1998).

In order to investigate the extent of ordination in microsatellite data attributable to population genetic structure, without using previous information on the origin of each individual, a Discriminant Analysis of Principal Components (DAPC, Jombart et al., 2010) was performed using Adegenet R package (Jombart et al., 2008). DAPC optimizes variation between clusters and minimizes variation within them, and it is free of assumptions such as Hardy-Weinberg and linkage equilibria (Jombart, Devillard & Balloux, 2010). At first data are transformed using a Principal Component Analysis (PCA), and then clusters are identified using discriminant analysis. The number of clusters (K) was identified by the *find.clusters* function using the “K-means” algorithm, and the Bayesian information criterion (BIC) was used to choose the most relevant K values for population structure. The discriminant analysis was then performed using the optimal number of principal components identified by a spline interpolation of the a-scores (i.e. the difference between the proportion of successful reassignment of the analysis and the values obtained using random groups).

The population genetic structure across the study area was also investigated using the Bayesian clustering algorithm implemented in TESS 2.3.1 and the geographical location of individuals as prior information (Chen, Durand, Forbes & François, 2007; Francois & Durand 2010). The analysis was performed by modelling admixture using a conditional autoregressive model (CAR). Preliminary analyses were carried out to assess model performance, with 20 000 steps (the first 5 000 were discarded as burn-in) and 10 replicates for each K value (i.e. the number of clusters) between 2 and 10. The final analysis contained 100 replicates for each K value, with K = 2–10; each run consisted of 80 000 steps, with the first 30 000 discarded as burn-in. The spatial interaction parameter was initially kept at the default value (0.6), and the updating option was activated. The model that best fitted the data was selected using the deviance information criterion (DIC). DIC values were averaged over the 100 replicates for each K value, and the most probable K value was selected as the one at which the average DIC reached a plateau. For the selected K value, the estimated admixture proportions of the 10 runs with the lowest DIC were averaged using CLUMPP 1.1.2 (Jakobsson & Rosemberg, 2007).

## Results

The final dataset consisted of a multi-locus genotype for 76 individuals at thirteen microsatellite loci, with 9.8% of missing data. Micro-Checker detected the possible occurrence of null alleles at locus *Cg14E1* in population 13 and at locus *Cg15F7* in population 11. Except for these two populations, no significant deviation from the Hardy-Weinberg and linkage equilibria was found after the Bonferroni correction was applied. Allelic richness and mean expected heterozygosity estimates for each population are shown in Table 1. Population 13 (Sila Massif, central Calabria) and population 4 (Foresta della Lama, Tuscan-Emilian Apennines) showed the highest values of genetic diversity, whereas the lowest values of heterozygosity and allelic richness were observed in population 12 (Gargano, N Apulia).

DAPC identified K = 5 as the best clustering option, being the one with the lowest BIC value. The optimization of the spline interpolation of the a-scores suggests to use only the first 11 principal components (accounting for 54,4% of the total variance) as the more informative ones for the discriminant analysis. The inspection of the scatterplot resulting by the DAPC analysis (Fig. 2) clearly identified five main genetic clusters, including individuals from: i) the Western Alps (pops. 1-2), ii) the Northern and Central Apennines (pops. 3-11), iii) the Gargano Promontory (pop. 12), iv) the Sila Plateau and Catena Costiera massif (central Calabria, pops. 13-14), and v) the Aspromonte Massif (southern Calabria, pop. 15).

**Figure 2.**
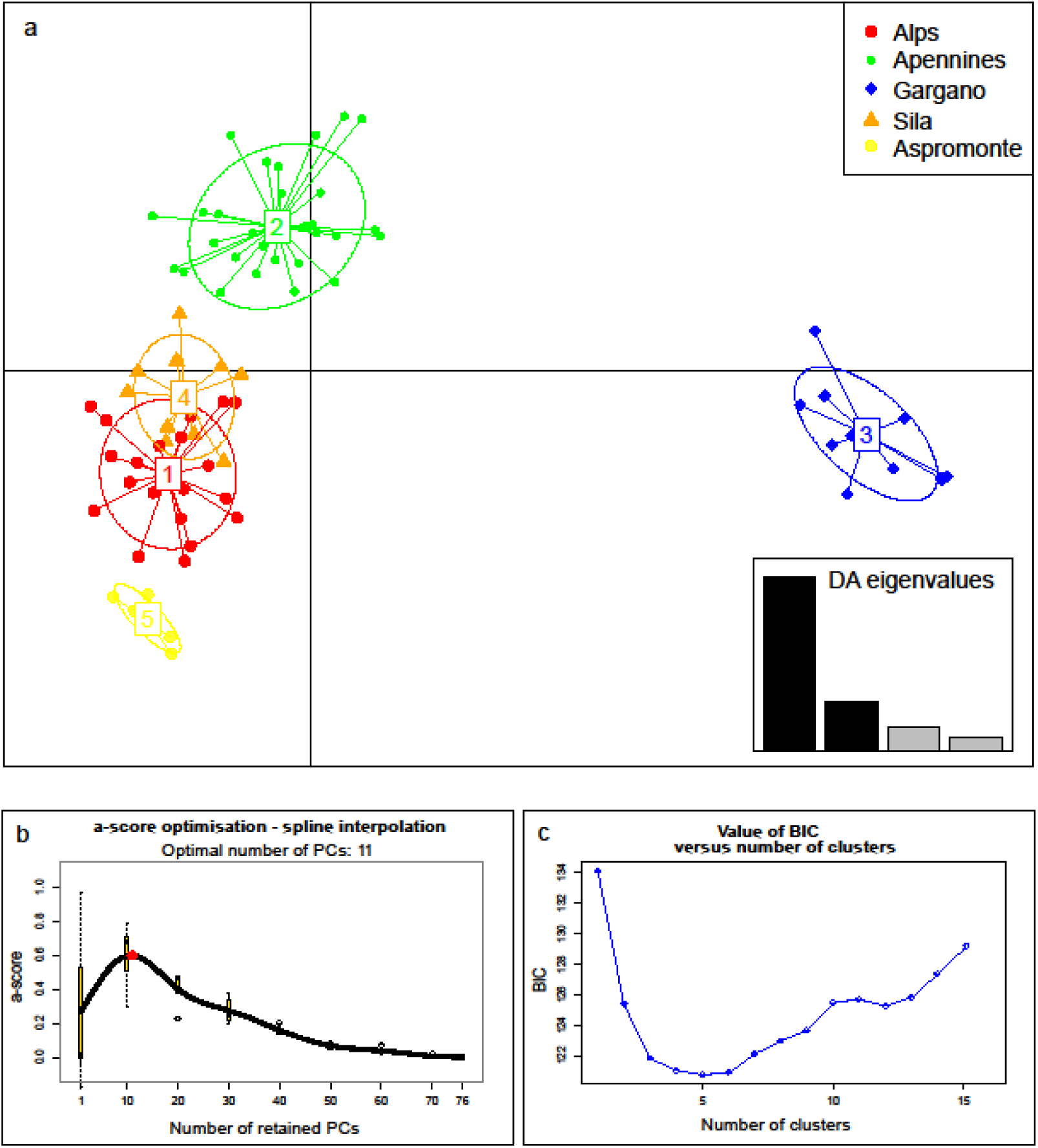
Discriminant Analysis of Principal Components (DAPC). (a) Scatterplot resulting from the DAPC performed on *Myodes glareolus* genotypes from the Italian peninsula, using K=5 as clustering option; axes represent the first two principle components; the box shows the relative contribution of the eigenvalues to the discriminant analysis; clusters are named following their geographic distribution. (b) Optimal number of informative principal components, suggested by the optimization of the spline interpolation of the a-scores. (c) line chart showing the BIC values versus the number of genetic clusters (K) ranging from 1 to 15.

The Bayesian clustering analyses carried out with TESS revealed a clear geographic structuring of genetic variation, consistent with results of the DAPC analysis. The plots of DIC values versus K values reached a plateau at K = 5 and only a minor decrease in the DIC values was observed at higher K values. The spatial distribution of the five clusters had a clear geographical structure: one is widespread in the Alps and, a lower frequency, in the Northern Apennines; one is found from Northern to Central Apennines; one is restricted to the Gargano Promontory region; one ranges from the Sila Plateau to the Catena Costiera, and one from the Catena Costiera to the Aspromonte Massif. Bar-plots showing the individual admixture proportions and pie-charts showing the average proportion of each cluster within each sampled population are given in Figure 3. Large genetic admixture is observed in individuals from Catena Costiera (pop. 14), as well as in those from northern (pop. 3-5) and central Apennines (pop. 6-11).

**Figure 3.**
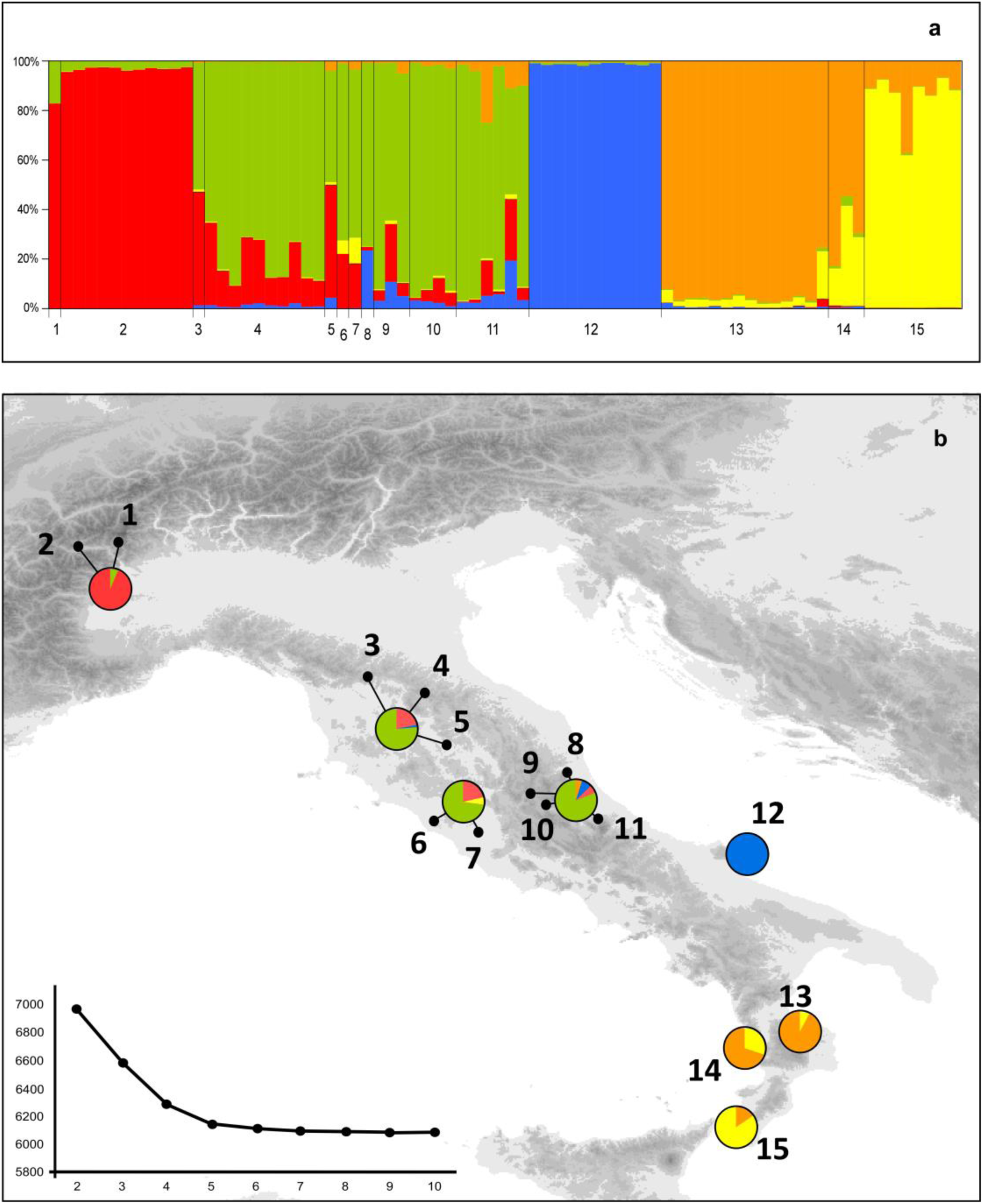
Genetic structure of Italian populations of *Myodes glareolus* at 13 microsatellite loci estimated using TESS. (**a**) The bar plot shows the admixture proportions of each individual for the five genetic clusters recovered. (**b**) The pie diagrams on the maps show the frequency distributions of each cluster among the populations; populations with only one individual were grouped with the nearest population as follow: (1,2), (3,4,5), (6,7), (8,9,10,11); the line chart shows the mean values of the DIC statistics (averaged over 100 runs) for the number of genetic clusters (K) ranging from 2 to 10. The map was drawn using the software Canvas 11 (ACD Systems of America, Inc.).

## Discussion

The geographic structure of microsatellite variation supports the existence of five main genetic clusters within *Myodes glareolus* in the Italian peninsula, in contrast to the four lineages identified by the mitochondrial DNA. Indeed, previous studies identified a genetic lineage from Europe widespread in northern Italy, a distinct genetic lineage in central Italy, a slightly differentiated lineage restricted to the Apulian region, and a highly differentiated lineage restricted to the Calabrian region (Colangelo et al., 2012; Filipi et al., 2015). Our data support for the independent evolution of these lineages, and identified further sub-structuring within the Calabrian region. We found evidence for genetic differentiation between populations from Aspromonte and Sila, and for admixture between these two groups within the Catena Costiera. Genetic differentiation is also corroborated by some morphological distinctiveness between these populations, which has led to the description of distinct subspecies: *Myodes glareolus curcio* (von Lehmann, 1961) for the Sila Massif and *Myodes glareolus hallucalis* (Thomas, 1906) for the Aspromonte Massif (Amori et al., 2008a; Viro & Niethammer, 1982). Moreover, our results suggest strong genetic isolation between southern populations and the other Apennine populations and claim for considering the Calabrian and Gargano lineages as independent evolutionary and conservation units, deserving special attention in conservation planning.

Strong genetic differentiation and high intra-specific variation of Calabrian populations is a fairly common pattern in both animal and plant species (Bisconti et al., 2018; Canestrelli et al., 2006; Canestrelli, Cimmaruta & Nascetti, 2008; Canestrelli, Aloise, Cecchetti & Nascetti 2010; Canestrelli, Sacco & Nascetti, 2012; Chiocchio, Bisconti, Zampiglia, Nascetti & Canestrelli, 2017; Hewitt, 2011; Vega, Amori, Aloise, Cellini, Loy & Searle, 2010). The Calabrian region is a well-known glacial refugium for temperate species in peninsular Italy, and provides one of the best examples of highly sub-structured refugia, a scenario known as refugia-within-refugia (Gomez & Lunt, 2007). Indeed, for most of the temperate species studied to date in this area, the Calabrian region provided suitable albeit fragmented habitats through most of Pleistocene, allowing long-term survival of relict populations (Bisconti et al., 2018; Senczuk, Colangelo, De Simone, Aloise & Castiglia, 2017). Accordingly, the Early-Middle Pleistocene origin of Calabrian bank vole populations was suggested by both molecular dating and fossil evidence (Colangelo et al., 2012; Sala & Masini 2007). Moreover, palynological data support the expansion of Alpine forests in Calabria during the Early-Middle Pleistocene transition, as a consequence of particularly humid glacial cycles (Capraro et al., 2005; Palombo, Raia & Giovinazzo, 2005). The southward expansion of forest habitats promoted southward colonization of several forest and woodland-dwelling species, and might have promoted the establishment of the bank vole populations in Calabria, which probably remained trapped after the following shrinking of woodlands. The almost complete absence of admixture between the Calabrian cluster and those located more to the north, strongly suggests an ancient isolation of Calabrian populations.

On the other hand, the sub-structuring of the bank vole populations within the Calabrian region appears of more recent origin and it is likely related to the high palaeogeographic instability of this region. According to palaeogeographic reconstructions, the repeated glacioeusthatic sea level oscillations of the Pleistocene caused repeated marine floods in the lowlands, turning the main mountain massifs into paleo-islands (Bonfiglio et al., 2002; Caloi, Malatesta & Palombo, 1989; Cucci, 2004; Ghisetti, 1979, 1981; Tansi, Muto, Critelli & Iovine, 2007; Tortorici, Monaco, Tansi & Cocina, 1995). The repeated insularization of Sila and Aspromonte massifs heavily affected population structure in most of the terrestrial fauna inhabiting these areas (Canestrelli et al., 2006, 2008, 2010, 2012) and could have triggered genetic differentiation also in the bank vole populations. The relatively high levels of genetic diversity and genetic admixture observed in these populations could be explained by a more recent secondary contact between the two gene pools. Under this scenario, the high level of genetic variation observed within the Calabrian Pleistocene refugia would be better explained by dynamic microevolutionary processes, which involve cycles of allopatric divergence and secondary contact, rather than by a prolonged demographic stability (Canestrelli et al., 2010).

Conversely, the population from the Gargano Promontory - described as a distinct subspecies *Myodes glareolus garganicus* (Hagen, 1958; see Amori et al., 2008a) - showed strong genetic divergence and low genetic diversity, probably as a consequence of a historical isolation and small population size, which favoured genetic erosion by drift. However, a role for the strong anthropogenic impact on the bank vole’s habitats, which affected this region during the last decades (Parise & Pascali, 2003; Ladisa, Todorovic, & Liuzzi, 2010), cannot be excluded.

Populations from Alps and from north-central Apennine belong to two different genetic clusters, in spite of substantial levels of admixture. This pattern is consistent with that showed by mtDNA data (Colangelo et al., 2012) and suggests good habitat connectivity and high levels of gene flow throughout Central Apennine and Alps, at least after the last glacial phase. Therefore, the genetic structure of bank vole populations in the Italian peninsula, as inferred by both microsatellites and mitochondrial DNA markers, is consistent with a scenario of independent evolution of multiple genetic lineages within distinct glacial refugia. Interestingly, the postglacial range expansion of the lineages from Northern and Central Italy was more extensive than that showed by the southern lineages, which appear still confined to their refugial areas. However, we suggest caution in considering the geographic and genetic isolation of Calabrian and Apulian populations. The paucity of observations from geographically intermediate populations, as well as the lack of morphological and genetic data does not allow to trace neither geographic nor genetic boundaries among these lineages. Further research should be focused on the intermediate areas, in order to ascertain the presence of bank vole populations and to estimate their genetic structure.

Our results have important implications for the management of the bank vole populations in Southern Italy. Due to its widespread distribution throughout most of Europe, *Myodes glareolus* is currently categorized as *Least Concern* by the IUCN red list of threatened species, both at the global (Amori et al., 2008b) and national (Rondinini, Battistoni, Peronace, & Teofli, 2013) level. Nevertheless, we identified at least three unique evolutionarily significant units (Moritz, 1994) in Southern Italy, with narrow and endemic ranges. Assessments of their demographic consistence, as well as of current threats to their populations will be compulsory in the near future, in order to ensure long-term conservation of these ancient and unique evolutionary lineages. This is particularly urgent considering that Italian laws do not protect vole species outside regional and national parks.

Concluding, this study highlights the importance of investigating species genetic structure with a multi-marker approach, in order to find hidden diversity and fine-scale genetic structuring also in supposedly well-known species. The analysis of bank vole genetic structure in the Italian peninsula revealed more biological diversity than expected, suggesting the need for thorough research also in other apparently well-known taxa. Finally, the evolutionary history of *Myodes glareolus* provides further evidence supporting the hypothesis that Pleistocene refugia were not so stable as previously thought, and that dynamic micro-evolutionary processes, triggered by the paleoclimatic and palaeogeographic instability of these areas, better explain the high levels of intraspecific diversity they harbour.

## Acknowledgements

We are grateful to Giuliano Milana for helping in sampling activities and to Paola Arduino for providing useful comments and suggestions on the manuscripts. This research was supported by grants from the Italian Ministry of Education, University and Research (PRIN project 2012FRHYRA), and from the Aspromonte National Park.

